# Acoustic Signatures of the Cerrado: Machine Learning Reveals Unique Soundscapes Across Diverse Phytogeographies

**DOI:** 10.1101/2024.01.12.575467

**Authors:** Bruno Daleffi da Silva, Linilson Padovese

## Abstract

This article explores the application of machine learning techniques in acoustic ecology to classify the formations of the Brazilian Cerrado (Forest, Savanna, and Grassland) based on their soundscapes. Considering the Cerrado’s importance for biodiversity and hydrology, as well as the challenges faced by the biome in the face of agricultural expansion, the study seeks more efficient and economical methods for identifying its physiognomies.

Five statistical models were developed and evaluated, using both traditional Machine Learning and Deep Learning, with the use of Mel-Frequency Cepstral Coefficients (MFCCs) and spectrogram images as input variables. The performance of these models was measured by accuracy, precision, and recall metrics, revealing a superiority of the Convolutional Neural Network (CNN), which, despite requiring greater computational cost and training time, provided high precision in the classifications and valuable insights through the application of the LIME explainability technique.

Moreover, the study proposes a majority vote classification methodology for frequently observed events, enabling reliable classifications through models with moderate performance. It is concluded that the choice of the ideal model for the classification of soundscapes of the Cerrado should consider a balance between accuracy, operational complexity, and efficiency. The conclusions of this study offer relevant directions for future research and the application of monitoring technologies in conservation and recovery efforts of biomes.

## 1. INTRODUCTION

The soundscape of the Cerrado, one of Brazil’s richest and most threatened biomes, has been the subject of intense research due to its biodiversity and ecological importance. This biome, covering about 23% of the national territory and spanning 12 states (IBGE-BDIA), is characterized by a semi-arid climate and dense shrub vegetation. However, the expansion of agricultural and livestock activities has led to a significant loss of its vegetation cover, with about 150,000 square kilometers of vegetation being deforested between 2000 and 2018 (IBGE, 2020, Annex 2). The degradation of the Cerrado not only threatens its flora and fauna, including endemic species, but also affects its critical role in the water and climate regulation of the country (DIAS, 1991).

Given this scenario, the restoration of degraded areas and the preservation of the existing biome become imperative. To this end, it is essential to understand the natural formation of the region, identify the loss of biodiversity, and determine the ideal formation for restoration, which may or may not correspond to the original biome, depending on the degree of degradation (CAVA, 2018). Traditionally, the process of identifying the phytophysiognomies of the Cerrado is based on qualitative analyses of flora and fauna, which require substantial field efforts (RIBEIRO apud EMBRAPA, 2008).

**Figure 1.**
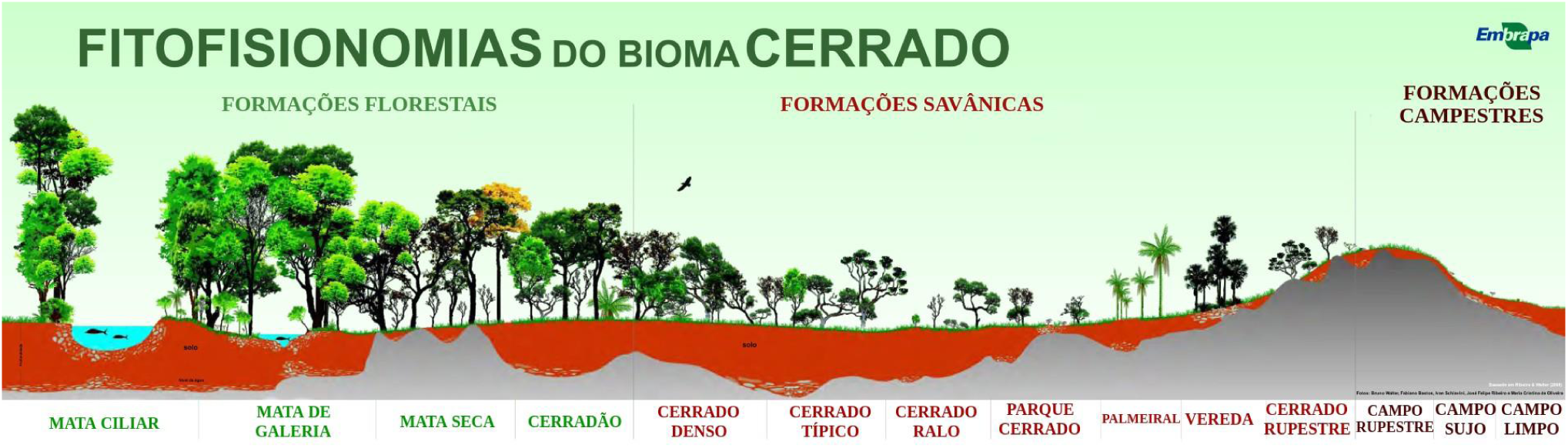
Phytophysiognomies of the Cerrado. Source: Embrapa

However, recent advances in Acoustic Ecology offer a promising alternative, using the soundscape to identify and classify environmental characteristics in an efficient and cost-effective way (Tucker, 2014).

This paper presents an innovative approach that integrates Artificial Intelligence in the analysis of the Cerrado’s soundscape, exploring the potential of machine learning models to discern its different natural formations. By using acoustic data from representative regions of the Forest, Savanna, and Grassland formations, this study aims to validate the hypothesis that similar ecological identities are reflected in similar acoustic landscapes. The efficiency in identifying and classifying these formations through their sound profiles is practical proof of the validity of this hypothesis.

The contribution of this work is threefold: first, it highlights the application of sound classification based on Artificial Intelligence; second, it considers the relationship between the complexity of the method and its practicality; and third, it emphasizes the use of LIME in spectrogram-based CNNs, a significant advancement that allows identifying the regions of the spectrum and frequency bands that characterize each formation. These advances not only enhance the precision of the analyses but also provide valuable insights for the conservation and restoration of the Cerrado.

## 2. OBJECTIVES

The core of this study was to evaluate the hypothesis that natural formations with similar ecological identities exhibit comparable acoustic landscapes. This evaluation was conducted through the development and comparison of machine learning models, which were applied to classify the natural formations of the Cerrado based on the soundscapes of the selected regions. The efficiency in identifying and classifying these formations through their sound profiles served as practical validation of this central hypothesis.

To achieve the main objective, the study was structured around the following guiding questions:

1. Which machine learning models, ranging from traditional statistical methods to advanced neural networks, are most efficient and effective in classifying the types of natural formations of the Cerrado based on their soundscapes?
2. How do the complexity and computational cost of the models impact the choice of the most appropriate method for classifying acoustic landscapes?
3. Which attributes of the sound signals are most relevant for classifying the acoustic landscapes of the Cerrado?

The proposed investigation sought to analyze the efficacy of models that use Mel Frequency Cepstral Coefficients (MFCCs) and spectrograms as input variables in the classification of acoustic landscapes. Machine Learning and Deep Learning models, such as Gradient Boosting, Random Forest, Logistic Regression, Multilayer Perceptron, and Convolutional Neural Networks, were employed.

Furthermore, the study examined the impact of the number of observations on the performance of the models and the importance of considering the simplicity of the method, the training and prediction response time, and the capacity to handle a large number of observations in a short period. Finally, this study aimed to understand the importance of the main characteristics of the spectrogram in classifying the acoustic landscapes of the Cerrado.

Therefore, the outlined objectives sought not only to validate the central hypothesis but also to contribute to the methodological advancement in the application of Artificial Intelligence for the analysis of soundscapes in ecological contexts.

## 3. LITERATURE REVIEW

The analysis of soundscapes, or acoustic landscapes, has emerged as a prominent approach in assessing biodiversity and understanding ecosystems. The soundscape consists of a complex web of sounds, known as biophony, geophony, and anthrophony, which reflect the interactions among living beings, natural phenomena, and human activities (Priestman, 2017). Biophony, in particular, is a vital indicator of the presence and behavior of species in an ecosystem, making it a valuable tool for biodiversity studies.

In the context of the Cerrado, a biome of extreme ecological importance and unique biodiversity, the use of soundscapes for environmental assessment is especially relevant. The Cerrado is a biodiversity hotspot, home to a wide variety of endemic species and crucial for maintaining important watersheds (DIAS, 1991). Acoustic analysis of the environment can provide detailed information about the ecological health of these areas, allowing for the detection of changes in species composition and ecosystem dynamics.

Recent studies have applied Machine Learning and Deep Learning methods to classify species and monitor biodiversity through sound recordings. For instance, Mehyadin et al. (2021) achieved 78% accuracy in bird classification using sound wave signals as a database. These advancements demonstrate the potential of combining sound landscape analysis techniques and Artificial Intelligence in the conservation and monitoring of biomes like the Cerrado.

The application of Machine Learning models, such as Gradient Boosting, has shown promising results in soundscape studies. Fonseca et al. (2017) demonstrated the effectiveness of this model, improving baseline performance by 8.2% in classifying acoustic scenes. Similarly, the use of Random Forest in audio signal classification systems achieved an overall correct classification rate of 99.25% in a study by Grama et al. (2017), highlighting this model’s ability to handle unbalanced datasets and identify sounds related to wildlife intruder detection.

Logistic Regression has also been successfully applied in the analysis of urban soundscapes, as shown by Noviyanti et al. (2019), who used MFCC coefficients to predict sound perception with Correct Classification Indices of up to 88.3%. Moreover, the Multilayer Perceptron was effective in multi-label classification of acoustic patterns in audio clips, as explored by Zhang et al. (2016), providing detailed insights into the distribution of various acoustic patterns in long-duration recordings.

Convolutional Neural Networks (CNNs) have stood out in soundscape analysis, particularly in the classification of spectrograms (Stowell et al., 2014; Pellegrini et al., 2020). Khamparia et al. (2019) explored the classification of environmental sounds using CNNs and achieved 77% accuracy on the ESC-10^1^ dataset. This study underscores the efficiency of CNNs in recognizing and classifying spectrogram images, which are visual representations of sound frequencies over time.

Acoustic Ecology, which originated with the pioneering works of Schafer (1977) and Truax (2001), has evolved with the development of Data Science and Artificial Intelligence. The integration of these disciplines has enabled a deeper and automated analysis of soundscapes, enhancing the understanding of ecological interactions and the impacts of human activities on the environment. The World Forum for Acoustic Ecology (WFAE), established in 1993, has been a platform for the dissemination and international recognition of this research area (Wrightson, 2000).

The application of these techniques in the Cerrado can be an effective strategy for preserving the biome. Identifying specific soundscapes associated with different vegetation formations in the Cerrado is a crucial step for implementing conservation and restoration measures. Acoustic analysis can assist in identifying degraded areas and evaluating the effectiveness of management practices, contributing to the sustainability and resilience of this ecosystem.

In summary, the literature review indicates that soundscape analysis, supported by advancements in Artificial Intelligence, is a promising approach for biodiversity assessment and ecosystem conservation. In the Cerrado, this approach can not only help understand the biome’s complexity but also guide preservation and recovery efforts in the face of increasing anthropogenic pressures.

## 4. METHODOLOGY

Adopting a structured methodical strategy, the study followed four crucial steps that mirror the data science cycle as proposed by Shearer (2000).

### 4.1. Data Collection

The collection of acoustic data was a crucial step in this study and was carried out with the assistance of high-precision equipment from the Laboratory of Acoustics and Environment (LACMAM) of the Polytechnic School of the University of São Paulo (USP). For a comprehensive analysis of the Cerrado soundscapes, three strategically located Ecological Stations (ESECs) were selected, each representing one of the main vegetation formations of the biome: forest, grassland, and savanna.

The chosen regions were Assis, representing the forest formation; Itirapina, as an example of the grassland formation; and Águas de Santa Bárbara, illustrating the savanna formation. These classifications were not assigned arbitrarily; they result from meticulous field studies conducted by experts who examined the characteristics and densities of the vegetation at each site. This initial classification was essential to establish a reference point for the subsequent acoustic analysis.

With the sound recording devices properly installed, data collection proceeded. The goal was to capture a broad spectrum of acoustic variations, including sounds emitted by fauna, noises generated by abiotic elements such as wind and rain, and any anthropogenic interferences. This holistic approach ensures that the sound profile captured is representative of the complexity and richness of the Cerrado biome.

The volume of data collected was substantial, amounting to approximately 3 TB of sound recordings covering the period from September to November. The audio data from the Itirapina ESEC were recorded in 5-minute files, with a sampling rate of 8 KHz and a depth of 16 bits. In contrast, the acoustic landscapes of the Santa Bárbara and Assis ESECs were recorded in 3-minute files, with a sampling rate of 32 KHz and a depth of 16 bits. Given the magnitude of the collected data, it was necessary to adopt a strategy to make the volume of data more manageable while maintaining the integrity and representativeness of the information.

For this purpose, a random selection of 7,500 files, totaling 70 GB of data, was made. This strategic reduction in the sample ensured a diverse representation of weather conditions and recording times, covering different scenarios such as winds, rains, and diurnal and nocturnal variations. This approach allowed the machine learning models to be trained efficiently, without compromising their ability to learn and recognize the patterns of the Cerrado soundscapes in various contexts.

The collection of acoustic data in three distinct regions of the Cerrado provides a solid foundation for subsequent analysis and machine learning modeling. By understanding the acoustic characteristics associated with each vegetation formation, this study aims to contribute significantly to the preservation and restoration of the Cerrado biome, as well as to the science of acoustic ecology as a whole.

### 4.2. Pre-Processing

After the collection of the sound data, the subsequent stage of the study involved meticulous pre-processing of the captured audios. From this phase onward, all analyses were carried out using the R programming software.

The first step in processing the information was the standardization of the duration of the recordings, restricting the analysis to the first minute of each sample. This uniformity was essential to ensure comparability among the recordings, regardless of their original duration.

Next, the sampling rates were resampled using the tuneR library in R. This function is inspired by the melfcc.m function, developed in the Matlab rastamat library (Hermansky, 1990 and Hermansky et al., 1994). This process was necessary to synchronize the datasets, which were recorded at different sampling rates across the ESECs. Resampling to 8KHz for the data from the Santa Bárbara and Assis ESECs ensured that all data were in a consistent format, facilitating subsequent analyses.

The Short-Time Fourier Transform (STFT) was applied using the spectro function of the seewave package. The seewave library, in turn, is based on the book “Animal Acoustic Communication: Sound Analysis and Research Methods” (Hopp et al. 1998).

The STFT is a technique that allows for the analysis of the frequencies of a signal over time, decomposing the sound signal into its frequency and amplitude components. The result was a three-dimensional database that captures the temporal evolution of frequencies and their respective amplitudes, providing a detailed representation of the acoustic content of the recordings.

The last step of the pre-processing involved normalizing the amplitudes, a crucial step to standardize the volume of the recordings and mitigate any distortions or variations in sound intensity that might affect the analysis, such as variations from recording equipment. Normalization was carried out to adjust the amplitude levels to a common scale, facilitating the comparison between different recordings.

At the end of this process, the audios were transformed into databases containing detailed information about the time, frequency, and amplitude of each of the Cerrado formations. This transformation created the necessary foundation for the supervised analyses that followed, allowing the machine learning models to be trained and validated with structured and standardized data, essential for the precise identification of the acoustic characteristics of each vegetation formation.

Finally, with the help of the ggplot2 library in R, the processed data were transformed into visual spectrograms. Spectrograms are graphical representations that illustrate how the frequencies of a signal vary over time, offering an intuitive and informative view of the studied sound environment. These images were fundamental for the modeling phase, enabling the application of machine learning techniques for the classification of the Cerrado’s acoustic landscapes.

Figure 2 displays examples of these spectrograms, illustrating the sonic diversity of the Cerrado formations.

**Figure 2.**
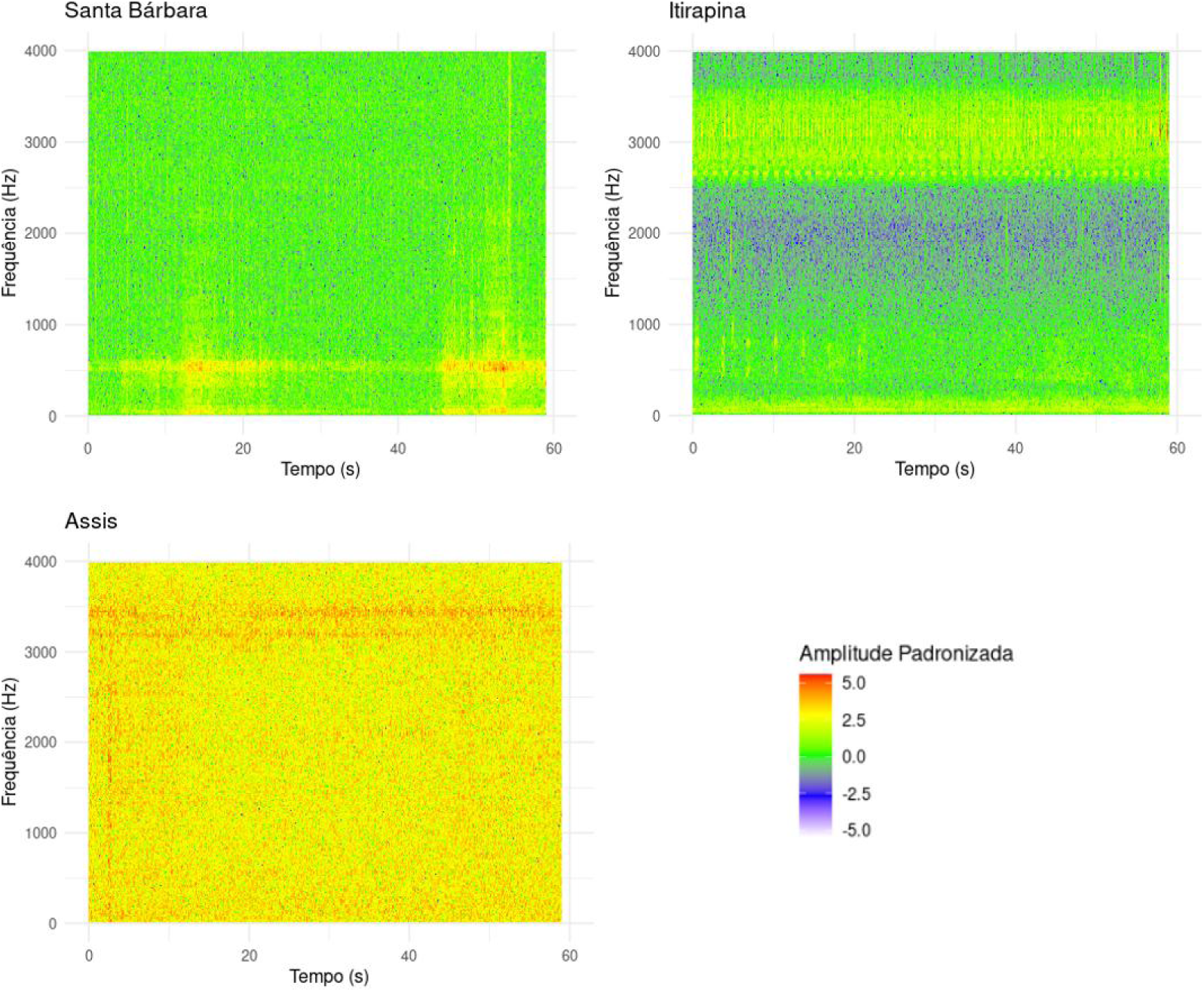
Example of spectrograms from the three regions/formations.

### 4.3. Modeling

The modeling phase was a crucial step in this study, where pre-processed information was processed through a range of advanced machine learning techniques. The selection of models was based on a thorough review of the literature and previous technological advancements, ranging from traditional statistical methods to deep neural networks, as described by Friedman (2001), Breiman (2001), Cox (1958), Rosenblatt (1958), and LeCun et al. (1998). For traditional models, such as Gradient Boosting, Random Forest, Logistic Regression, and Multilayer Perceptron, Mel Frequency Cepstral Coefficients (MFCCs) (DAVIS et al., 1980) were used as input variables. On the other hand, for the Convolutional Neural Network (CNN), spectrogram images were used as the basis for classification.

The machine learning models Gradient Boosting, Random Forest, Logistic Regression, and Multilayer Perceptron (MLP) were trained using MFCCs as explanatory (input) variables. Except for the MLP, the models were implemented with the R tidymodels library, which provides an integrated and consistent approach to statistical modeling and machine learning (KUHN, 2022), following the principles of tidyverse.

Among the packages offered by tidymodels are “tune” and “parsnip”. The “tune” package plays a fundamental role in optimizing the hyperparameters of machine learning models. Hyperparameters are model settings that need to be specified before training and are not learned from the data. They can influence model effectiveness, and their optimization allows the evaluation and comparison of different models, improving their performance.

On the other hand, the “parsnip” package is used for model specification. It provides a consistent interface for defining and configuring machine learning models, regardless of the underlying computational engine that implements it. This package was extensively used in this study to define and adjust the hyperparameters of the Gradient Boosting, Random Forest, and Logistic Regression models.

For the Gradient Boosting model, using the boost_tree function from the parsnip package, the following hyperparameters were adjusted: mtry, min_n, tree_depth, sample_size, learn_rate, and loss_reduction. Here, mtry refers to the number of variables available for splitting at each node of the tree, min_n is the minimum number of observations in the nodes, tree_depth is the maximum depth of any tree, sample_size is the fraction of data used to build each tree, learn_rate is the learning rate, and loss_reduction refers to the minimum loss reduction required to make a new split in the tree.

The Random Forest model, implemented through the rand_forest function of the parsnip package, had the hyperparameters mtry and min_n adjusted. Mtry, as in the previous model, is the number of variables available for splitting at each tree node, while min_n is the minimum number of observations in the nodes.

In Logistic Regression, using the multinom_reg function from the parsnip package, the hyperparameters penalty and mixture were adjusted. The penalty refers to the amount of regularization applied, which helps to avoid overfitting, while mixture determines the type of regularization applied (L1, L2, or a mix of both).

The implementation of CNN and MLP was carried out using the TensorFlow (GOOGLE, 2023) and Keras (CHOLLET et al., 2023) libraries, which are open-source tools for machine learning and deep neural networks. These libraries offer a flexible and powerful environment for building, training, and validating complex models, such as CNNs, which are particularly suited for processing visual and acoustic data.

The MLP architecture was developed based on the Keras and Tensorflow packages of R (derived from Python). This architecture featured six intermediate layers, including two dense layers with 30 and 18 neurons, respectively, plus a dropout layer and a normalization layer, arranged between the dense layers. The total number of parameters adjusted in the MLP model was 1,101.

The architecture of the CNN was composed of several layers, including convolutional layers, batch normalization layers, max pooling layers, and spatial dropout layers. The network starts with an input layer with dimensions of 250×250×3, representing the dimensions of the input images.

Specifically, the model has three convolutional blocks, each with two convolutional layers, followed by a batch normalization layer, a max pooling layer, and a spatial dropout layer. The number of filters in the convolutional layers progressively increases in each block, starting with 32, then 64, and finally 128. All convolutional layers use ReLU activation functions and have a kernel size of 5×5, with ‘same’ padding to ensure the output has the same dimension as the input.

After the three convolutional blocks, the network has a global average pooling layer, followed by a flatten layer and a dense layer with 128 units and ReLU activation function. A dropout layer with a rate of 0.4 is applied before the output layer, which has three units and a softmax activation function, corresponding to the three classification categories.

The model has a total of 815,139 adjustable parameters, of which 814,243 are trainable and 896 are non-trainable. During training, the model was compiled using the Adam optimizer with a learning rate of 0.001, loss function ‘categorical_crossentropy’, and evaluation metric ‘accuracy’. Two callback functions were used: one to reduce the learning rate when validation loss stopped improving (callback_reduce_lr_on_plateau) and another to stop training when the validation accuracy exceeded 98% after 10 consecutive epochs above 97%.

The model was trained for up to 300 epochs, with a batch size of 64. Validation data was used to monitor model performance and adjust the learning rate as needed. The use of callbacks allowed for more efficient training, avoiding overfitting and saving computational time.

The modeling methodology also included a strategic division of data, separating the total set into three distinct parts: training, validation, and blind test. The training base was used to adjust the models and teach them the patterns of the data. The validation base played the role of optimizing hyperparameters and preventing overfitting, ensuring the generalization of models. The blind test base, composed of data not exposed to the models during training, was crucial to test the effectiveness of the models under new and unknown conditions, providing a reliable assessment of their performance.

Each model underwent a careful training and validation process, aiming to maximize its accuracy and generalization capacity. Accuracy, precision, and recall metrics were used to evaluate the performance of each algorithm, providing a complete view of the effectiveness of the models. Additionally, explainability techniques, such as LIME, were planned to be applied to the CNN, with the aim of interpreting the decisions made by the model and identifying the most relevant acoustic features for the classification of the Cerrado’s acoustic landscapes.

In summary, modeling was carried out with methodological rigor that allowed not only efficient classification of acoustic landscapes but also a detailed analysis of the models, contributing to the advancement of knowledge at the intersection between Acoustic Ecology and Data Science.

### 4.4. Explainability

Understanding the decisions made by machine learning models, especially in complex neural networks like CNNs, is essential to validate the reliability and applicability of the obtained results. In this context, the analysis of explainability focused on the predictions generated by the CNN, employing the technique of LIME (Local Interpretable Model-agnostic Explanations) as proposed by Ribeiro et al. (2016). LIME is a methodology that facilitates the interpretation of complex models by introducing perturbations in the input data and observing variations in the model’s predictions. This technique induces the CNN to formulate local linear models that are intrinsically more comprehensible, uncovering the impact of acoustic attributes on the classification of soundscapes.

The methodology implemented by LIME involves generating modified versions of the input data and the subsequent analysis of changes in the CNN’s predictions. This analysis allows identifying which features of the data are decisive for the model’s decisions. By applying LIME to specific spectrograms, it is possible to identify which frequency segments or temporal patterns are considered most relevant by the CNN in classifying a soundscape as belonging to a specific formation of the Cerrado.

The adopted explainability approach not only elucidated the relevance of acoustic attributes but also promoted greater transparency in the model’s predictive decisions. With an understanding of the factors influencing the CNN’s predictions, a more solid basis is obtained for applying the results in conservation and management initiatives of the Cerrado biome.

In summary, the methodology employed in this study allowed for the development of effective statistical models and provided a detailed understanding of predictive decisions. The use of LIME, in particular, ensured a consistent and reliable approach, enabling the validation of the CNN’s predictions and a deeper interpretation of the acoustic patterns characteristic of the various formations of the Cerrado. Clarity in the decisions of the models is a crucial aspect for the progress of the application of Artificial Intelligence in ecological and environmental research.

## 5. RESULTS

The application of machine learning models for the classification of the Cerrado’s acoustic landscapes yielded significant results. The performance of each model was evaluated based on accuracy, precision, and recall metrics, as presented in Table 1.

**Table 1.** Performance of the developed models according to the analyzed metrics.

| Model | Accuracy | Precision | Recall | Training time |
| --- | --- | --- | --- | --- |
| Gradient Boosting | 93% | 93% | 93% | 5,86 minutes |
| Random Forest | 92% | 92% | 92% | 5,31 minutes |
| Logistic Regression | 82% | 83% | 82% | 9,89 seconds |
| Multilayer Perceptron | 83% | 79% | 69% | 2,33 minutes |
| CNN | 99% | 98% | 98% | ~ 8 days |

The Convolutional Neural Network (CNN) demonstrated superiority, achieving 99% accuracy, 98% precision, and 98% recall. This remarkable performance indicates that the CNN was capable of identifying the different formations of the Cerrado from the soundscapes with high precision. However, the substantially longer training time (∼8 days) compared to the other models should be considered in the context of practical application.

Nevertheless, the ‘repetitive classification’ strategy proved to be an efficient technique when using models with moderate performance for frequently occurring events. This strategy involves repeatedly applying the same model to multiple small samples of the same sound event, and the final decision is made based on the most frequent response among the classifications, a method known as majority voting. This approach derives its effectiveness from consistency and convergence evidenced in accurately identifying the Cerrado’s natural formation across numerous observations, echoing the principles of the Law of Large Numbers (Bernoulli, 1713).

In any case, although the CNN requires significant investment in terms of time and computational resources, it brings an additional advantage: the ability to explain its decisions. With the help of LIME (Ribeiro et al. in 2016), a model interpretation technique, we were able to identify which sound features are most important for classifying the different formations of the Cerrado. This is clearly demonstrated in Figure 3, where LIME helps to clarify which aspects of the audio recordings are most relevant for the CNN, providing not just an accurate classification but also a deeper understanding of the acoustic data we are analyzing.

**Figure 3.**
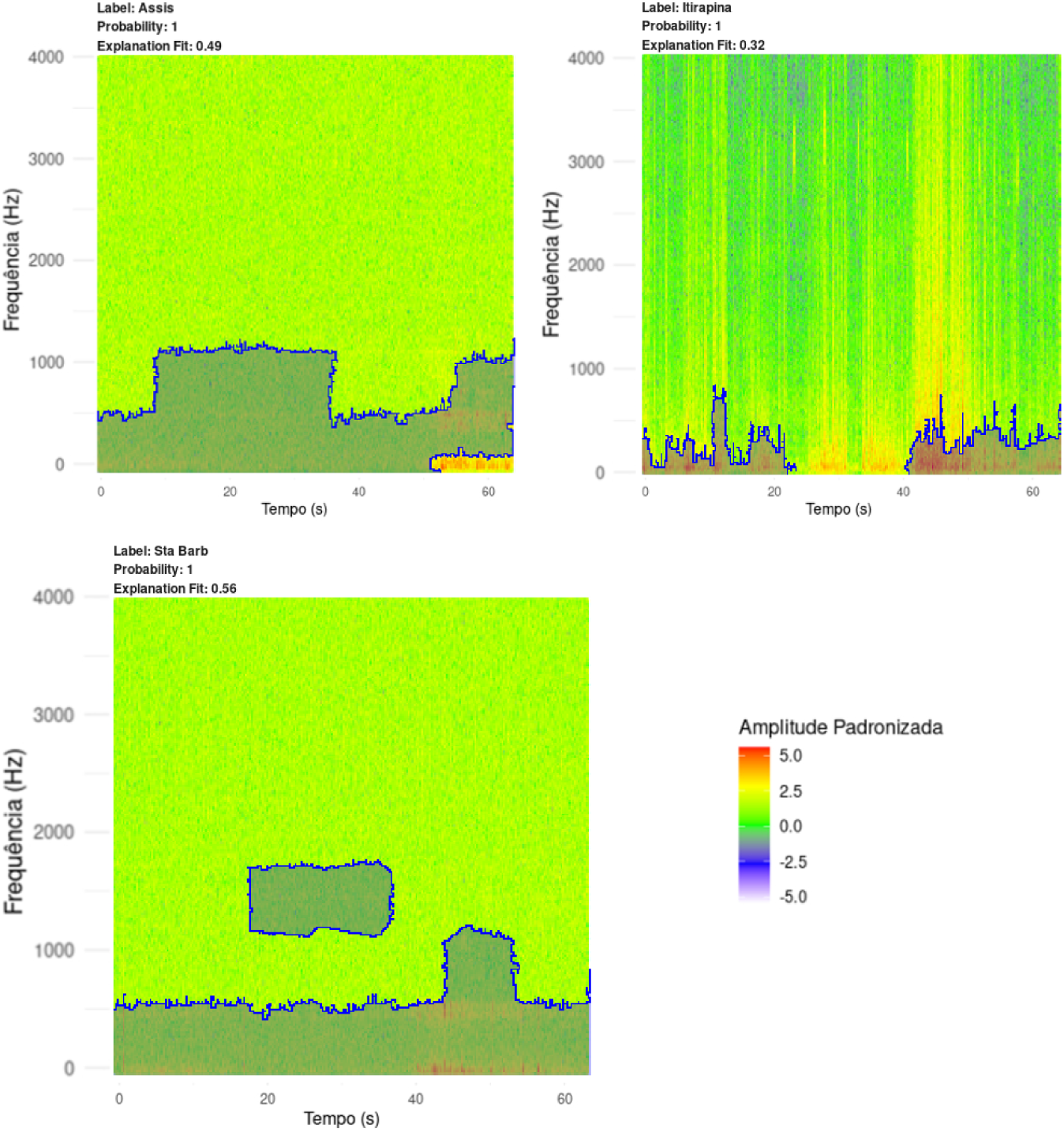
Application of LIME to Explain the CNN on Examples of Spectrograms.

## 6. DISCUSSIONS

The advancements achieved in this study underscore the synergy between Acoustic Ecology and Data Science in the effective classification of soundscapes, providing valuable analytical tools to advance the understanding and preservation of the Cerrado. The explored models exhibited variabilities in performance, but the applicability and efficiency of their implementation proved to be equally important factors for their selection in environmental analysis projects.

The remarkable performance of the CNN in quantitative results reflects its aptitude for extracting and learning complex features from soundscapes. However, it is important to note that the high accuracy of this model may be offset by the simplicity and agility of less sophisticated methods, such as Gradient Boosting or Random Forest, which prove advantageous in various practical scenarios, especially when frequent evaluations are required.

The methodological approach for classifying high-exposure events played a crucial influence, suggesting that reasonably performing models, when applied iteratively, can provide accurate classification and converge to the true identification of the natural formation. This technique relies on the consistency of results generated when dynamizing the analysis of multiple samples, establishing a reliable precedent for environmental categorizations.

The use of LIME in analyzing the CNN provided additional insights, allowing not only the identification of the Cerrado formations with high precision but also an understanding of the specific acoustic properties that determine each classification. Such elucidation highlights the importance of combining different techniques and models to extract the maximum value from the analyzed data.

In conclusion, the research emphasized the relevance of balancing various factors in choosing models for classifying acoustic landscapes. The multiparametric perspective of this article underscores the efficacy of interdisciplinarity in the application of artificial intelligence for the monitoring and conservation of natural ecosystems.

## 7. CONCLUSION

The investigation conducted in this study provided robust evidence in support of the central hypothesis that similar natural formations have similar soundscapes. Through the application of machine learning and Deep Learning models, it was possible to classify the formations of the Cerrado with high precision based on their acoustic landscapes, validating the hypothesis and demonstrating the potential of these technologies in ecological analysis.

In response to the first guiding question, it was identified that both traditional statistical methods and advanced neural networks are efficient in classifying acoustic landscapes. However, the Convolutional Neural Network (CNN) stood out as the most effective model, albeit requiring significant training time and computational resources.

Regarding the second question, the complexity and computational cost of the models influenced the choice of the most appropriate method. While the CNN offered the best performance, models such as Gradient Boosting and Random Forest proved to be viable and efficient alternatives, especially when considering the repetitive classification strategy and the principle of majority voting for high-exposure events.

As for the third question, the attributes of the sound signals that proved most relevant for classifying the Cerrado’s acoustic landscapes were the specific frequencies identified by the models, especially by the CNN, whose interpretation was enhanced by the use of LIME.

This study concludes that the integration of Artificial Intelligence in the analysis of soundscapes is a promising approach for biodiversity assessment and ecosystem conservation. The models developed and tested offer valuable tools for identifying natural formations, contributing to the preservation and recovery of threatened biomes like the Cerrado. The choice of the appropriate model should be guided by a balance between accuracy, efficiency, and practicality, considering the specific needs of each application. The explainability of the models, particularly the CNN, reinforces confidence in classification decisions and provides valuable insights for future research and practical applications.

## Footnotes

1 The ESC-10 dataset consists of 10 classes of sounds, while the ESC-50 contains 50 different classes of environmental sounds, such as beaches, rain, bird songs, etc. (Piczak, 2015).

